# Development and optimization of a diluted whole blood ELISpot assay to test immune function

**DOI:** 10.1101/2024.01.20.576465

**Authors:** Ricardo F. Ungaro, Julie Xu, Tamara A. Kucaba, Mahil Rao, Scott C. Brakenridge, Philip A. Efron, Robert W. Gould, Richard S. Hotchkiss, Monty B. Mazer, Patrick W. McGonagill, Lyle L. Moldawer, Kenneth E. Remy, Isaiah R. Turnbull, Charles C. Caldwell, Vladimir P. Badovinac, Thomas S. Griffith

**Author notes:** Address reprint requests to Thomas S. Griffith, Ph.D., Department of Urology, University of Minnesota, 3-125 CCRB, 2231 6^th^ St. SE, Minneapolis, MN 55455.

## Abstract

**Background:** Sepsis remains a leading cause of death worldwide with no proven immunomodulatory therapies. Stratifying Patient Immune Endotypes in Sepsis (‘SPIES’) is a prospective, multicenter observational study testing the utility of ELISpot as a functional bioassay specifically measuring cytokine-producing cells after stimulation to identify the immunosuppressed endotype, predict clinical outcomes in septic patients, and test potential immune stimulants for clinical development. Most ELISpot protocols call for the isolation of PBMC prior to their inclusion in the assay. In contrast, we developed a diluted whole blood (DWB) ELISpot protocol that has been validated across multiple laboratories.

**Methods:** Heparinized whole blood was collected from healthy donors and septic patients and tested under different stimulation conditions to evaluate the impact of blood dilution, stimulant concentration, blood storage, and length of stimulation on *ex vivo* IFNγ and TNFα production as measured by ELISpot.

**Results:** We demonstrate a dynamic range of whole blood dilutions that give a robust *ex vivo* cytokine response to stimuli. Additionally, a wide range of stimulant concentrations can be utilized to induce cytokine production. Further modifications demonstrate anticoagulated whole blood can be stored up to 24 hours at room temperature without losing significant functionality. Finally, we show *ex vivo* stimulation can be as brief as 4 hours allowing for a substantial decrease in processing time.

**Conclusions:** The data demonstrate the feasibility of using ELISpot to measure the functional capacity of cells within DWB under a variety of stimulation conditions to inform clinicians on the extent of immune dysregulation in septic patients.

## INTRODUCTION

Sepsis remains a leading cause of mortality in the United States^1,2^. Although sepsis survival has significantly improved over the past decade, sepsis incidence rates continue to increase due (in part) to an aging population with greater comorbidities, making sepsis an ongoing public health challenge^3,4^. Improved acute sepsis treatment protocols have led to increased sepsis survivorship; however, up to 50% of sepsis patients never fully recover and go on to develop chronic critical illness (CCI)^5^. CCI is characterized by persistent immune dysfunction manifested as recurrent infections, sepsis recidivism, and poor long-term outcomes^6^. Identifying interventions to improve immune function in sepsis patients is central to improving outcomes for patients with CCI. The first step toward immunomodulatory therapies for sepsis is to define and characterize immune dysfunction in sepsis patients.

Current immunophenotyping efforts in sepsis rely on static metrics of immune function such as soluble cytokines, cell surface marker expression, or cellular transcriptomics^5,7–9^. These efforts have been largely unsuccessful, in part because they often fail to directly assess immune function. Further, these existing techniques are time intensive, limiting the practical application of their findings to direct patient care. The overarching hypothesis of the Stratifying Patient Immune Endotypes in Sepsis (“SPIES”) consortium holds that serial ELISpot assays measuring *ex vivo* IFNγ or TNFα production by peripheral blood cells will be superior to other static tests of host immunity because ELISpot quantifies immune cell function rather than phenotype^10^.

ELISpot is an *ex vivo* assay used to study the responsiveness of the adaptive or innate immune cells. Initially developed to enumerate antibody-secreting cells, it is most used to detect cytokine-secreting cells^11,12^. ELISpot is attractive as a sepsis diagnostic/prognostic tool because the ELISpot platform is already used for FDA-approved clinical testing. Current commercially deployed ELISpot assays rely on the isolation of peripheral blood mononuclear cells (PBMC) from whole blood by density gradient centrifugation. Processing the blood in this manner removes mature granulocytes, platelets, and plasma components that may influence the function of the lymphocytes and monocytes^13–15^. Moreover, the steps required for PBMC isolation introduce the potential for cell loss and/or sample contamination. In response to these challenges, we previously established a novel ELISpot approach using undiluted whole blood and found that septic patients with suppressed T cells IFNγ production had an increased risk of adverse outcomes^16^. These data suggest whole blood ELISpot as a candidate assay to characterize immune function in critically ill patients.

In this study we sought to further develop the whole blood ELISpot assay as a candidate diagnostic and prognostic test for patients with sepsis. It is important to recognize that multiple factors can influence the performance of the ELISpot assay, including the number of cells used in each well, source and subset composition of the cells analyzed, dose and type of stimulant used, condition of sample prior to testing (specifically, temperature and length of storage), and duration of *ex vivo* stimulation. Thus, the data presented herein outline our methodology for developing a diluted whole blood (DWB) ELISpot assay using the production of two important cytokines (i.e., IFNγ and TNFα) for defining immune responsiveness by sepsis patients.

## MATERIALS and METHODS

### Patient populations and blood collection

Blood samples were collected from septic patients within the first three days of ICU admission in sodium heparin blood collection tubes (Becton Dickinson #367878, Franklin Lakes, NJ), as part of a multi-center, prospective observational study. In most cases, the blood was used in ELISpot within 1 hour of collection; however, some studies stored the blood at either 4°C or room temperature for 24 hours before use. Sepsis was defined according to Sepsis-3 criteria^17^, and inclusion criteria consisted of ICU admission from the emergency department or operating room for community-acquired sepsis, transfer to the ICU from an inpatient ward for the development of hospitalized sepsis, or the development of sepsis in a previously uninfected ICU patient. All patients were managed under standardized clinical management protocols. A central Institutional Review Board (IRB) approval from the sponsoring institution (University of Florida) was obtained, followed by concurrence from the IRB at each site. Informed consent was obtained from each patient or their surrogate decision-maker. Heparinized blood samples were also collected from healthy volunteer control subjects recruited using agreed upon criteria.

### Immune cell function using enzyme-linked immunosorbent spot (ELISpot) assay

ELISpot assays were conducted using human IFNγ or TNFα Immunospot^®^ kits (CTL Inc., Cleveland, OH) with several important modifications. Heparinized whole blood was diluted with kit media, and 50 µL of the diluted whole blood was added to each well. Duplicate samples were incubated in wells that contained either media alone, 500 ng/mL anti-CD3 (clone HIT3a: #300332, BioLegend, San Diego, CA) and 5 µg/mL anti-CD28 (clone CD28.2: #302934, BioLegend) mAb, or 2.5 ng/mL LPS (from *E. coli*, Serotype O55:B5: #ALX-581-013-L002, ENZO Life Sciences, Farmingdale, NY). Samples were incubated for 4 h or 22 h (+/- 1 h) at 37° C with 5% CO_2_. Wells were washed twice with PBS and twice with PBS-0.05% Tween-20 until the residual RBC layer was adequately removed. Biotinylated anti-human-IFNγ or -TNFα detection mAb (provided with each kit) was then added to each well for 2 h at room temperature. After washing thrice with PBS-0.05% Tween-20, a streptavidin-alkaline phosphatase conjugate was then added to each well for 30 minutes at room temperature. Finally, plates were washed twice with PBS-0.05% Tween-20 and twice with distilled water before adding the blue developer substrate solution for 15 minutes. Plates were washed with water at least 3 times to stop the reaction and allowed to air dry for at least 1 h before analysis. ELISpot plates were quantitated using a CTL S6 Entry ELISpot reader. Results are presented as the number of spot-forming units (SFU), as determined using the Immunospot^®^ SC software suite (version 7.0.30.4).

### Statistical Analysis

Statistical analyses were performed with the GraphPad Prism v10 (GraphPad Software Inc., San Diego, CA) for Mac OS X software package. Statistical comparisons of two groups were done using the unpaired nonparametric Mann-Whitney test. Statistical comparisons of more than two groups were done using Kruskal-Wallis tests, where the multiple comparisons were corrected with Dunn’s post hoc test. Statistical details for each experiment can be found in the figure legends.

## RESULTS

### Diluting whole blood improves spot recognition

The concentration of numerous cytokines, chemokines, and other metabolites in the blood increase during a septic event^18^. These soluble mediators drive the canonical sepsis-induced systemic inflammatory response, but also effect the function of immune cells in the blood^19–21^. To assess the cytokine-producing ability of the circulating adaptive and innate immune cells within the context of the entire host cellular and plasma milieu, we developed a novel ELISpot protocol to accommodate the use of whole blood during the stimulation step, incorporating all soluble and cellular blood components in the assay. Previously developed protocols for ELISpot using PBMC call for ∼2.5X10^5^ cells per assay; however, the ideal number of whole blood cells needed for a whole blood assay was unknown. We reasoned undiluted whole blood would overload the well with cells – especially erythrocytes – and restrict the detection of the cells producing cytokine upon stimulation. Further, the effect of other soluble and cellular components, such as mature granulocytes, on stimulated cytokine production was unknown. Thus, we first measured the effect of serial dilution of whole blood on assay performance. Heparinized whole blood from healthy donors was first diluted 1:2, 1:5, 1:10, 1:20, 1:50, 1:100, 1:200, and 1:500 with kit media before adding to the wells of an ELISpot plate. It is important to note these dilutions were of the initial blood sample, but all samples are diluted an additional 1:4 in the assay well (50 µL diluted blood plus 150 µL assay medium). Thus, the final blood dilutions were ultimately 1:8, 1:20, 1:40, 1:80, 1:200, 1:400, 1:800, and 1:2000. IFNγ production was measured in anti-CD3/CD28 mAb-stimulated wells; TNFχξ production was measured after LPS stimulation. The number of spot-forming units (SFU) were highest when the blood was diluted 1:5 before stimulation, and then progressively decreased when the blood was diluted further (Figure 1). Less diluted blood (1:2) resulted in a lower number of SFU, presumably because the higher concentration of erythrocytes interfered with the ability of the white blood cells to contact the well membrane. Based on these data, we chose to dilute the blood 1:10 with kit media before adding it to the ELISpot plate for stimulation in the subsequent assays.

**Figure 1.**
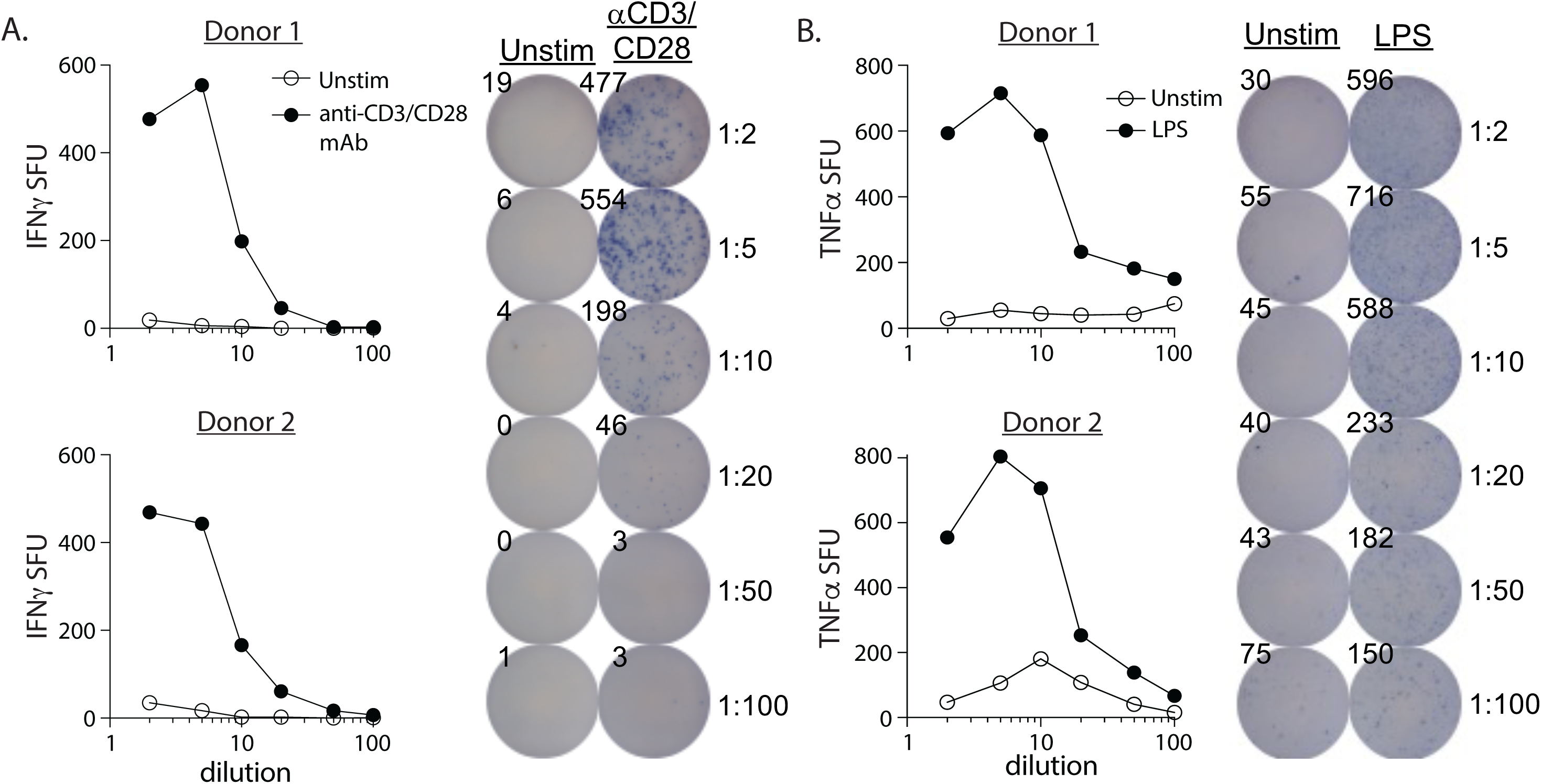
Titration of diluted whole blood from healthy donors show the dynamic range for detecting the IFNγ or TNFα response after 22 hours stimulation with anti-CD3/CD28 mAb or LPS, respectively. Whole blood was diluted 1:2, 1:5, 1:10, 1:20, 1:50, 1:100, 1:200, and 1:500 with kit media prior to adding 50 µL to the wells of an ELISpot plate for 22 hours stimulation with either anti-CD3/CD28 mAb (0.5 µg/mL and 5 µg/mL) or LPS (2.5 ng/mL). Unstimulated blood samples were step up in parallel wells to determine spontaneous IFNγ or TNFα SFU numbers. The number of (A) IFNγ and (B) TNFα spot-forming units (SFU) are shown from 2 healthy donors. Representative well images showing SFU are presented below each stimulation condition.

### IFNγ production in ELISpot is more dependent on the concentration of anti-CD3 mAb than anti-CD28 mAb

Physiological antigen-specific T cell activation requires T cell receptor recognition of cognate peptide:MHC complexes (‘signal 1’) along with CD28 ligation of CD80/CD86 (‘signal 2’) on the antigen presenting cell^22^. *Ex vivo* incubation of T cells with anti-CD3ε and anti-CD28 mAb can similarly activate the T cells. This stimulation is antigen-independent, causing polyclonal T cell activation. We next examined the impact of the presence and concentration of the anti-CD3 and anti-CD28 mAb on the production of IFNγ in the ELISpot assay. 1:10 DWB was stimulated for 22 hours with varying concentrations of either anti-CD3 or anti-CD28 mAb. The number of IFNγ SFU decreased with each 1:2 dilution of the anti-CD3 mAb while anti-CD28 mAb concentration was held constant (Figure 2A). By comparison, the number of IFNγ SFU was unchanged when the anti-CD3 mAb was held constant in the setting of decreasing anti-CD28 mAb concentrations (Figure 2B). These results highlight the importance of both CD3 and CD28 engagement for stimulating IFNγ in the DWB ELISpot assay.

**Figure 2.**
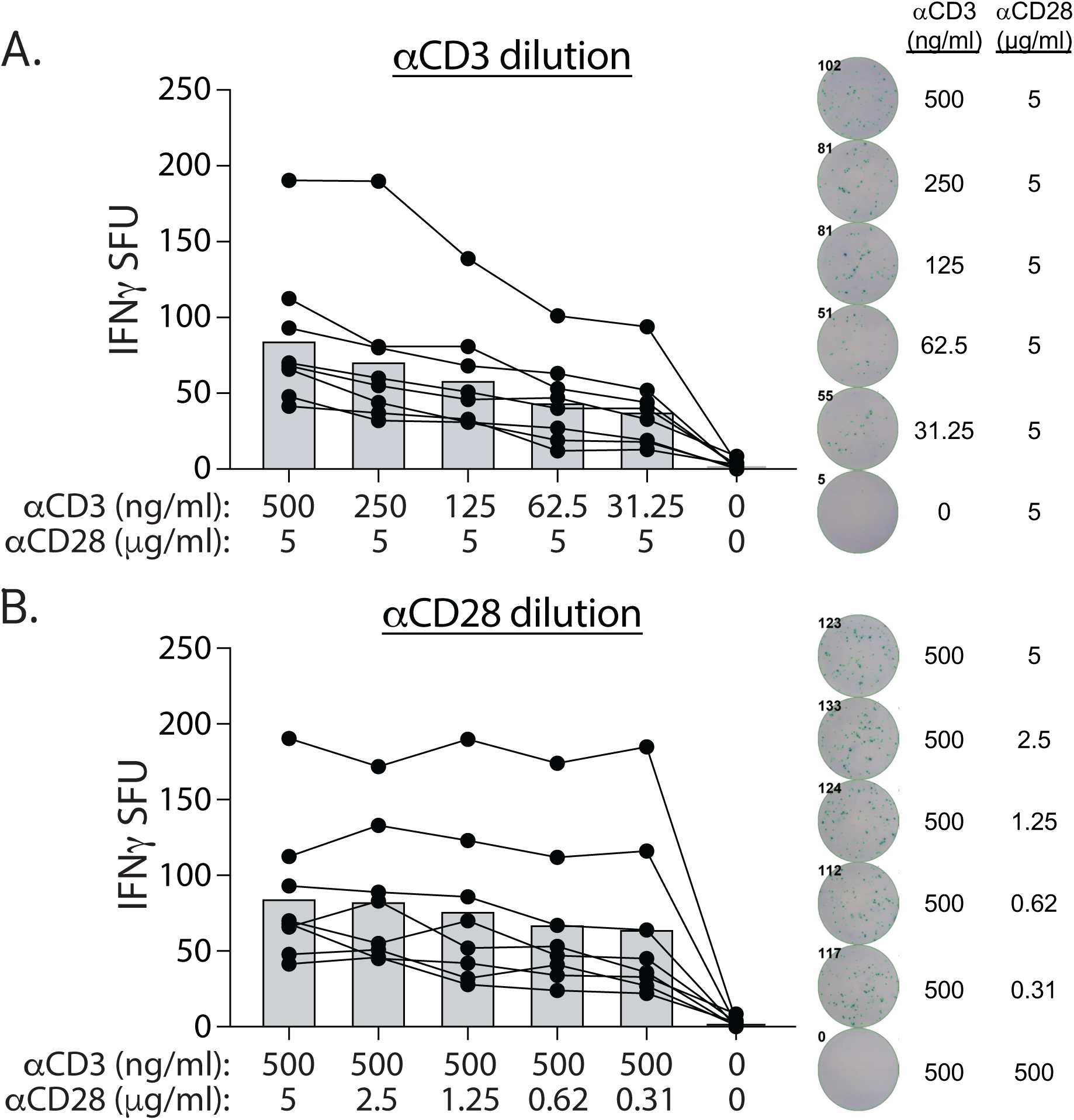
The response to anti-CD3/CD28 mAb stimulation is dependent on the dose of anti-CD3 mAb used. Whole blood was diluted 1:10 with kit media prior to adding 50 µL to the wells of an ELISpot plate for 22 hours stimulation with varying concentrations of anti-CD3 mAb and/or anti-CD28 mAb (as indicated in figure). The number of IFNγ spot-forming units (SFU) are shown from 7 healthy donors. The number of IFNγ SFU decreased in a dose-dependent manner when the concentration of anti-CD3 mAb decreased (A), but not when the anti-CD28 mAb concentration decreased (B). Lines in each graph connect samples from the same donor stimulated under the different conditions. Representative well images showing SFU are presented next to each stimulation condition.

### Blood can be stored for up to 24 hours at room temperature without losing significant activity in ELISpot

Timely processing of the blood for is important to generate accurate and actionable clinical data. Practically, the logistics of clinical laboratory testing mean it is not always feasible to process a patient blood sample immediately after it is drawn. Moreover, there is a growing trend in sending blood samples to centralized laboratories for analysis, which can mean a significant delay between phlebotomy and assay performance. Here, we tested the stability of the adaptive and innate immune response in the DWB ELISpot assay and sought to determine the best conditions for short term blood storage to minimize any loss of ELISpot functionality. Blood was collected from healthy donors and sepsis patients and then either assayed immediately (‘fresh’) or stored at 4°C or room temperature for 24 hours before running on ELISpot. The only statistically significant effect of storage (compared to fresh blood) was seen when blood from healthy donors was stored at 4°C for 24 hours before stimulation (Figure 3A). Cold storage for 24 hours repressed both anti-CD3/CD28 mAb-induced IFNγ and LPS-induced TNFα. There was a slight decline in number of SFU if the blood was held for 24 hours at room temperature, but this was an insignificant decrease when compared to fresh blood. Similar trends toward decreased SFU occurred when sepsis patient blood was stored at 4°C or room temperature for 24 hours, but these decreases were not statistically significant as compared to fresh patient blood (Figure 3B). These results suggest the window of time in which the samples need to be run on the ELISpot can be extended up to 24 hours after phlebotomy and demonstrate room temperature is the optimal storage condition.

**Figure 3.**
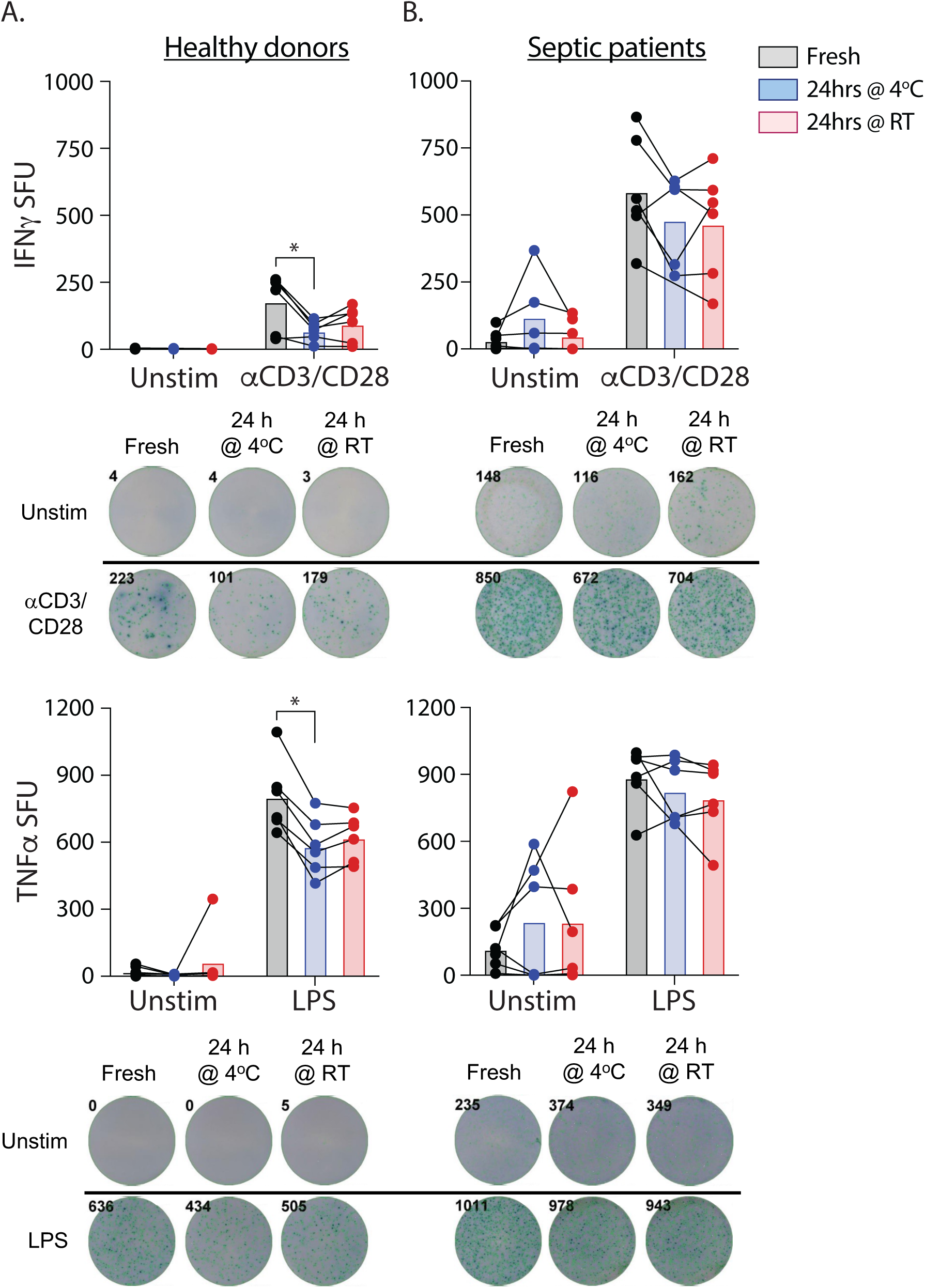
Blood can be stored for up to 24 hours without losing significant activity in ELISpot. Blood samples from 6 healthy donors (A) or 6 sepsis patients (B) were used immediately (‘fresh’) in the ELISpot assay or stored at 4°C or room temperature for 24 h before stimulation. Samples were diluted 1:10 with kit media before adding 50 µL to the wells of an ELISpot plate. The diluted whole blood was unstimulated or stimulated with either anti-CD3/CD28 mAb (0.5 µg/mL and 5 µg/mL) or LPS (2.5 ng/mL) for 22 hours to determine the number of IFNγ and TNFα spot forming units (SFU), respectively. In each graph, samples from the same patient are connected by the line. * p ≤ 0.05 using Kruskal-Wallis tests, where the multiple comparisons were corrected with Dunn’s post hoc test. Representative well images showing SFU are presented below each stimulation condition.

### 4-hour stimulation with anti-CD3/CD28 mAb or LPS is sufficient to induce IFNγ and TNFα production in the DWB ELISpot assay

Current PBMC-based ELISpot protocols call for a stimulation period of 20-24 hours^23^. Combined with the time needed for assay setup and the post-stimulation assay processing, this means results may not be available for up to 48 hours after phlebotomy. One of the long-term goals of the SPIES consortium is to use the ELISpot platform to identify immunosuppressed sepsis patients who would benefit from immunotherapy. The clinical imperative of prompt treatment requires getting results back to the bedside swiftly, enabling the initiation of the appropriate therapy in a timely manner. Thus, we were interested to see how shortening the stimulation period from 20-24 hours to only 4 hours would impact assay resolution. We chose a 4-hour stimulation because we reasoned this would allow information to become available within the same day as the blood draw. Even though there were significantly more IFNγ SFU after 22-hour stimulation with anti-CD3/CD28 mAb than seen after stimulation for 4 hours, there was still a robust signal in the 4-hour stimulation wells compared to unstimulated wells (Figure 4A). By comparison, there was no difference in number of TNFα SFU when the LPS stimulation lasted 4 or 22 hours (Figure 4B). The similarity in the LPS response is likely due to the release of preformed TNFα contained in the innate cells (i.e., monocytes and neutrophils) in the blood. Importantly, these results suggest ELISpot stimulation can potentially be truncated to permit for a more rapid completion of the entire protocol to reveal the results within 24 hours after blood draw.

**Figure 4.**
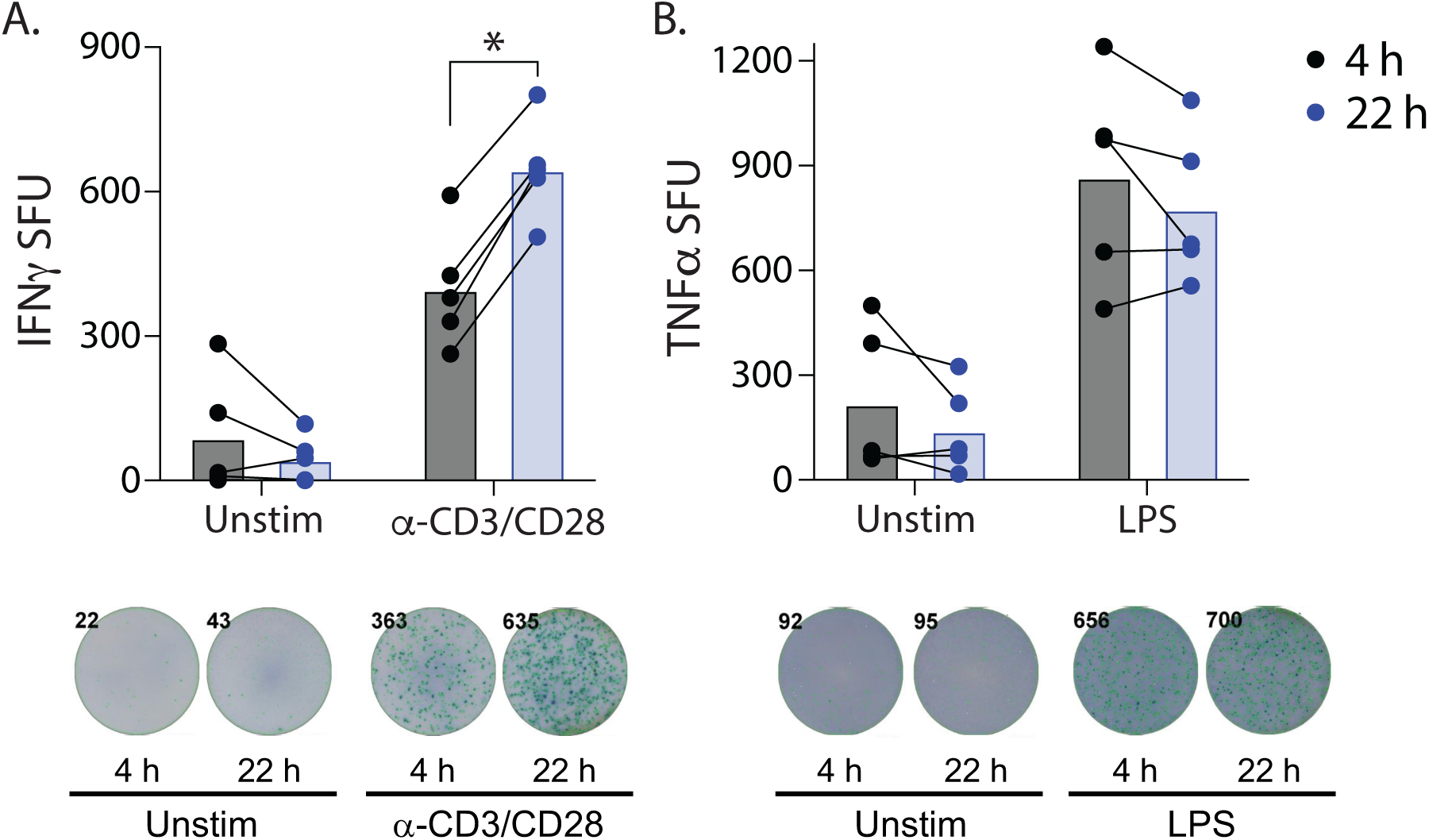
Reducing the ELISpot stimulation time to 4 hours maintains the ability to measure IFNγ and TNFα spot forming units. Blood samples from 5 septic patients were diluted 1:10 with kit media before stimulation with either anti-CD3/CD28 mAb (0.5 µg/mL and 5 µg/mL) or LPS (2.5 ng/mL) for either 4 or 22 hours to determine the number of (A) IFNγ and (B) TNFα spot forming units (SFU). In each graph, samples from the same patient are connected by the line. * p ≤ 0.05 using unpaired nonparametric Mann-Whitney test.

**Figure 5.**
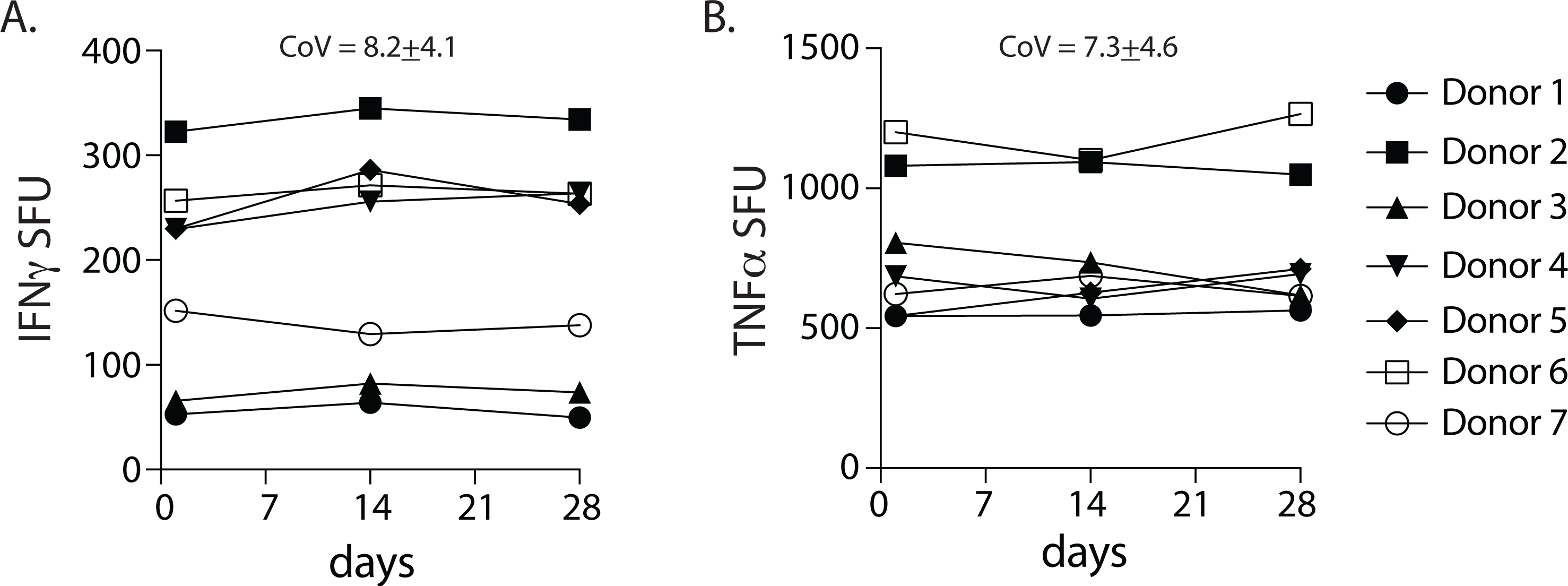
Optimization of ELISpot assay conditions leads to little variance in results from same healthy donors over time. Three blood samples from taken from 7 different healthy donors at 2-week intervals. Blood was diluted 1:10 with kit media before stimulation with either anti-CD3/CD28 mAb (0.5 µg/mL and 5 µg/mL) or LPS (2.5 ng/mL) for 22 hours to determine the number of IFNγ and TNFα spot forming units (SFU), respectively. In each graph, samples from the same patient are connected by the line.

### Serial blood draws from healthy donors demonstrate reproducibility of DWB ELISpot

One goal for the refinements to the DWB ELISpot protocol was to develop a highly reproducible assay to measure immune function in sepsis patients for use at each of the SPIES study sites. To verify assay consistency, we performed three serial blood draws from healthy donors spaced two weeks apart and determined the IFNγ and TNFα response after stimulation with anti-CD3/CD28 mAb or LPS for 22 hours. The number of IFNγ and TNFα SFU for each donor was quite consistent with each subsequent test, where the average coefficient of variance for the IFNγ and TNFα were 8.2±3.8 and 7.3±4.3, respectively.

## DISCUSSION

One of the defining hallmarks of sepsis is the development of a dysfunctional immune system that can either be proinflammatory, coagulopathic, and/or immune suppressed depending upon the source and site of infection, patient age and comorbidities, and timing since infection^1,3–6,17^. Several parameters have been used to define host immunity in the sepsis patient, but these often rely on static biomarkers that measure the concentration of a protein or transcript, or the phenotype of a cell^5,7–9^. The underlying goal of immunotyping sepsis patients is to understand the ability of the patients to respond to an infectious challenge. These static measures do not assess the ability of the immune system to participate in an effective immune response to a pathogen and instead reflect the response of the immune system to previous infection or tissue damage.

It has been nearly 50 years since the first report of a whole blood assay for measuring human immune functional responses^24^, but few of these assays have been translated into an actionable clinical test. The ELISpot assay is an attractive tool to characterize immune function in critically ill patients as it measures the ability of immune cells to respond to stimulation. Importantly, the ELISpot platform has been used clinically and ELISpot protocols and instrumentation are widely available. ELISpot is the foundational assay used in the SPIES consortium to interrogate the cytokine-producing capacity of both adaptive and innate immune cells within the blood of sepsis patients after stimulation with different agonists. We recently demonstrated that those sepsis patients who were more severely immune suppressed – using ELISpot as the function bioassay – had an increased risk of late mortality^10^. The purpose of this report was to delineate the methodology used to optimize the DWB ELISpot protocol so it could be employed in a multi-institutional research study and provide background for future clinical use. This report also reveals where the standard ELISpot protocol can be modified, based on the needs of the investigator to facilitate greater utility and action of the results.

ELISpot measures the number of cytokine-secreting cells at the single-cell level after *ex vivo* stimulation. Provided the requisite capture/detection mAb pairs are available, one strength of ELISpot is ability to detect cells producing nearly any cytokine of interest. We focused our assessment of the functional state of adaptive immunity by evaluating anti-CD3/CD28-stimulated T cell production of IFNγ, while innate immunity was tested by LPS-stimulated monocyte production of TNFα. Future studies will expand our interrogation of sepsis-induced immune dysfunction by measuring the number of cells producing cytokines like IL-6 or IL-10. Another benefit of ELISpot over other functional assays (e.g., intracellular cytokine staining for flow cytometry) is the ability to detect as few as one cytokine secreting cell in 100,000 cells per well^23^. Many protocols call for the use of PBMC in the ELISpot assay, but density gradient centrifugation separates the PBMC from granulocytes, platelets, and plasma proteins. Each of these components has the potential to influence the function of adaptive and innate immune cells in the blood. Using whole blood in the ELISpot allows for a more faithful representation of the blood as multicellular tissue, but we found it necessary to dilute the blood was necessary before adding it to the assay plate. The main reason for this was to prevent overloading the well with cells – especially erythrocytes – which may prevent the formation of a single cell layer on the membrane. We have also found that higher numbers of erythrocytes in the well can confound the results because of the possibility of releasing alkaline phosphatase via hemolysis^25^, increasing background signals within the assay well.

It is important to note that the use of (diluted) whole blood in the ELISpot also reduces the amount of sample processing needed before adding the sample to the assay plate. Depending on the type of study being performed, the assay can be performed either on-site or at centralized facility. Our data show no significant loss in cytokine-producing ability of the leukocytes when the blood was kept at room temperature for as long as 24 hours after collection. These results further suggest the potential of shipping samples from multiple locations to one site for processing, especially in cases where a site may not have sufficient personnel or equipment to perform the ELISpot assay.

We were intrigued by the data showing how the stimulation time of the ELISpot can be reduced to 4 hours and still detect a significant number of cytokine-producing T cells and monocytes. One obvious benefit of reducing the stimulation time is the ability to obtain information on the cytokine-producing capacity of immune cells in the blood of sepsis patients more quickly (potentially the same day as the blood draw) than with the more traditional overnight (20-24 hour) stimulation period. Rapidly providing data on immune dysregulation to the clinician may impact the course of treatment for the sepsis patient. Such a shorter stimulation time may prove to be beneficial when blood samples are processed at a centralized facility. It is also formally possible that different stimulation times can teach the researcher something about the function of different immune cell subsets. For example, a 4-hour stimulation with anti-CD3/CD28 mAb most likely induces IFNγ production from memory T cells, which are programmed to respond quickly during a secondary response^26–28^. Stimulation for 22 hours, by comparison, will encompass both the rapid response of memory T cells along with the slower response by naïve T cells. In contrast, innate immune cells respond rapidly after TLR ligation^29^, providing a potential explanation for no difference in the number of TNFα SFU when LPS was used as the stimulus in our DWB assay (Figure 4).

In summary, the results presented herein represent current methodology for developing a standardized DWB ELISpot protocol to conduct a prospective, multi-center observational study to immunologically endotype sepsis patients with the goal to predict clinical outcomes. The key parameters optimized included the amount of blood used in each well, dose and type of stimulant used, impact of sample storage (i.e., temperature and time) prior to use, and length of stimulation. Flexibility regarding adaptive and innate stimuli used and cytokine measured can reveal important information about different immune cell populations within the blood. It is likely we will modify future iterations of the ELISpot assay to address new questions arise in our investigation of the underlying mechanisms that impact sepsis patient outcomes.

## Notes

This work was supported in part by the National Institutes of Health grants: GM-132364 (including a supplement; to L.L.M), GM-142481 (to S.C.B.), GM-140806 (to P.A.E.), GM-126928 (to R.S.H.), GM-133756 (to I.R.T.), GM-134880 (to V.P.B.), and GM-1480881 (to T.S.G.). V.P.B. is a University of Iowa Distinguished Scholar. T.S.G. is the recipient of a Research Career Scientist award (IK6BX006192) from the Department of Veterans Affairs.

Conflict of Interest: M.B.M., K.E.R., and I.R. T. are members of Immune Functional Diagnostics, LLC (IFDx LLC) and receive no direct financial compensation. IFDx LLC is developing predictive metrics in critical illness and this technology is evaluated in this research. S.C.B, L.L.M., R.S.H., and the University of Florida may receive royalty income based on a technology developed by S.C.B. and others and licensed by Washington University in St. Louis to IFDx LLC. That technology is evaluated in this research. C.C.C. and the University of Cincinnati may receive royalty income based on a technology developed by C.C.C. and others and licensed by Washington University in St. Louis to IFDx LLC. That technology is evaluated in this research.

## REFERENCES

1. Rhee C, Dantes R, Epstein L, et al. Incidence and Trends of Sepsis in US Hospitals Using Clinical vs Claims Data, 2009-2014. JAMA. 2017;318(13):1241–1249.

2. Rudd KE, Johnson SC, Agesa KM, et al. Global, regional, and national sepsis incidence and mortality, 1990-2017: analysis for the Global Burden of Disease Study. Lancet. 2020;395(10219):200–211.

3. Vincent JL, Marshall JC, Namendys-Silva SA, et al. Assessment of the worldwide burden of critical illness: the intensive care over nations (ICON) audit. Lancet Respir Med. 2014;2(5):380–386.

4. Esper AM, Moss M, Lewis CA, Nisbet R, Mannino DM, Martin GS. The role of infection and comorbidity: Factors that influence disparities in sepsis. Crit Care Med. 2006;34(10):2576–2582.

5. Rincon JC, Efron PA, Moldawer LL. Immunopathology of chronic critical illness in sepsis survivors: Role of abnormal myelopoiesis. J Leukoc Biol. 2022;112(6):1525–1534.

6. Kahn JM, Le T, Angus DC, et al. The epidemiology of chronic critical illness in the United States*. Crit Care Med. 2015;43(2):282–287.

7. Balch JA, Chen UI, Liesenfeld O, et al. Defining critical illness using immunological endotypes in patients with and without sepsis: a cohort study. Crit Care. 2023;27(1):292.

8. Fenner BP, Darden DB, Kelly LS, et al. Immunological Endotyping of Chronic Critical Illness After Severe Sepsis. Front Med (Lausanne). 2020;7:616694.

9. Scicluna BP, van Vught LA, Zwinderman AH, et al. Classification of patients with sepsis according to blood genomic endotype: a prospective cohort study. Lancet Respir Med. 2017;5(10):816–826.

10. Barrios EL, Mazer MB, McGonagill PW, et al. Adverse outcomes and an immune suppressed endotype in sepsis patients with reduced interferon-gamma ELISpot. JCI Insight. 2023.

11. Czerkinsky CC, Nilsson LA, Nygren H, Ouchterlony O, Tarkowski A. A solid-phase enzyme-linked immunospot (ELISPOT) assay for enumeration of specific antibody-secreting cells. J Immunol Methods. 1983;65(1-2):109–121.

12. Hutchings PR, Cambridge G, Tite JP, Meager T, Cooke A. The detection and enumeration of cytokine-secreting cells in mice and man and the clinical application of these assays. J Immunol Methods. 1989;120(1):1–8.

13. De Groote D, Zangerle PF, Gevaert Y, et al. Direct stimulation of cytokines (IL-1 beta, TNF-alpha, IL-6, IL-2, IFN-gamma and GM-CSF) in whole blood. I. Comparison with isolated PBMC stimulation. Cytokine. 1992;4(3):239–248.

14. English D, Andersen BR. Single-step separation of red blood cells. Granulocytes and mononuclear leukocytes on discontinuous density gradients of Ficoll-Hypaque. J Immunol Methods. 1974;5(3):249–252.

15. Kirchner H, Kleinicke C, Digel W. A whole-blood technique for testing production of human interferon by leukocytes. J Immunol Methods. 1982;48(2):213–219.

16. Mazer MB, C CC, Hanson J, et al. A Whole Blood Enzyme-Linked Immunospot Assay for Functional Immune Endotyping of Septic Patients. J Immunol. 2021;206(1):23–36.

17. Singer M, Deutschman CS, Seymour CW, et al. The Third International Consensus Definitions for Sepsis and Septic Shock (Sepsis-3). JAMA. 2016;315(8):801–810.

18. Hotchkiss RS, Coopersmith CM, McDunn JE, Ferguson TA. The sepsis seesaw: tilting toward immunosuppression. Nat Med. 2009;15(5):496–497.

19. Heidarian M, Griffith TS, Badovinac VP. Sepsis-induced changes in differentiation, maintenance, and function of memory CD8 T cell subsets. Front Immunol. 2023;14:1130009.

20. Martin MD, Badovinac VP, Griffith TS. CD4 T Cell Responses and the Sepsis-Induced Immunoparalysis State. Front Immunol. 2020;11:1364.

21. Silva EE, Skon-Hegg C, Badovinac VP, Griffith TS. The Calm after the Storm: Implications of Sepsis Immunoparalysis on Host Immunity. J Immunol. 2023;211(5):711–719.

22. Jenkins MK, Johnson JG. Molecules involved in T-cell costimulation. Curr Opin Immunol. 1993;5(3):361–367.

23. Lehmann PV, Zhang W. Unique strengths of ELISPOT for T cell diagnostics. Methods Mol Biol. 2012;792:3–23.

24. Eskola J, Soppi E, Viljanen M, Ruuskanen O. A new micromethod for lymphocyte stimulation using whole blood. Immunol Commun. 1975;4(4):297–307.

25. Koseoglu M, Hur A, Atay A, Cuhadar S. Effects of hemolysis interferences on routine biochemistry parameters. Biochem Med (Zagreb). 2011;21(1):79–85.

26. Berard M, Tough DF. Qualitative differences between naive and memory T cells. Immunology. 2002;106(2):127–138.

27. Pennock ND, White JT, Cross EW, Cheney EE, Tamburini BA, Kedl RM. T cell responses: naive to memory and everything in between. Adv Physiol Educ. 2013;37(4):273–283.

28. Surh CD, Sprent J. Homeostasis of naive and memory T cells. Immunity. 2008;29(6):848–862.

29. Kawasaki T, Kawai T. Toll-like receptor signaling pathways. Front Immunol. 2014;5:461.

